# Semi-field evaluations of three botanically derived repellents against the blacklegged tick, *Ixodes scapularis* (Acari: Ixodidae)

**DOI:** 10.1101/2022.01.12.476114

**Authors:** Xia Lee, Colin Wong, Joel Coats, Susan Paskewitz

## Abstract

Three compounds derived from botanicals sources, ethyl perillyl carbonate, geranyl isovalerate, and citronellyl cyclobutane carboxylate, were tested for repellent activity against *Ixodes scapularis* Say in a semi-field trial. Tick drags were treated with the compounds or with N, N-diethyl-m-toluamide (DEET) at high (0.25mg/cm^2^) or low (0.15mg/cm^2^) concentrations. Negative controls included untreated drags and drags treated with acetone, the carrier for all repellents. Freshly treated drags (within 20 minutes) were used to collect *I. scapularis* ticks at a county park in Wisconsin. To assess effectiveness, we measured tick encounter rates, detachment rate, and time to detachment. None of the repellent treatments resulted in significantly fewer encounters compared to both control treatments. However, the percentage of ticks that detached within 3 min was significantly higher on drags treated with repellents compared to controls. DEET was the most effective, repelling 69.7 - 87% of ticks by 3 min, but the effectiveness of the three test compounds was still high, ranging from 42% to 87% of ticks detaching by 3 min. For time to detachment, there were no significant differences between DEET and the three test compounds. We conclude that these botanically-derived repellents were effective against *I. scapularis* in a semi-field trial and could be viable alternatives to DEET.

---

In the United States, the incidence of vector-borne diseases has been increasing with tick-borne diseases (TBDs) accounting for greater than 75% of reported cases (Rosenberg et al. 2018). The blacklegged tick, *Ixodes scapularis* Say, and the western blacklegged tick, *Ixodes pacificus* Cooley & Kohls, are responsible for the transmission of the main etiological agent of Lyme disease, *Borrelia burgdorferi*, which accounts for greater than 80% of all cases of TBDs (Eisen et al. 2016, Rosenberg et al. 2018, Lehane et al. 2021). Other ticks such as the American dog tick, *Dermacentor variabilis* Say, and the lone star tick, *Amblyomma americanum* L., are important vectors responsible for the transmission of the causative agents of Rocky Mountain spotted fever (*Rickettsia rickettsia*) and ehrlichiosis (*Ehrlichia chaffeensis* and *E. ewingii*), respectively (Mixson et al. 2006, Hecht et al. 2019). Management strategies aiming to reduce the burden of TBDs have relied on a toolbox of approaches that primarily focus on reducing human and tick encounter rates through landscape manipulation, use of broad spectrum acaricide, increased signage in tick infested areas, and personal prevention (Eisen and Dolan 2016, Lee et al. 2020). Recommendations for personal prevention include frequent tick checks, wearing permethrin-treated clothing and proper application of tick repellents on skin or clothes while outdoors (CDC, 2021).

Currently, N, N-diethyl-m-toluamide (DEET), a synthetic compound, is the most prominent active ingredient in repellents on the market and is available in concentrations ranging from 5-100%. Within the EPA’s database of 630 registered repellents against ticks, products with DEET as the active ingredient make up 80% of the market (EPA, 2021). Despite decades of use with a strong safety record, the use of DEET remains controversial due to public concerns over potential toxicological effects (Veltri et al. 1994, Osimitz and Grothaus 1995, Diaz 2016, Swale and Bloomquist 2019, Haleem et al. 2020). In addition, DEET is one of the most commonly detected organic contaminants in water and its impact on the environment has yet to be fully assessed (Merel and Synder 2016, Dos Santos et al. 2019).

Recent interest in plant-based natural products with low environmental impact has stimulated a growth in studies of botanical essential oils (EOs) as potential repellents for ticks (Jordan et al. 2012, Meng et al. 2016, Kim et al. 2021). Cedarwood, citronella, rosemary, geranium, nutmeg, clove, garlic and peppermint oil are a few promising EO’s that have been shown to be as effective as DEET in laboratory assays (Carroll et al. 2011, Meng et al. 2016, Benelli and Pavela 2018). Despite their reported effectiveness, it’s difficult to bring EO’s to market as a repellent due to their high volatility, which limits the period of efficacy against ticks. They are also chemically complex and composition of an EO can vary depending on methods of production. Nonetheless, EO’s are valuable sources for deriving new compounds with potential repellent or acaricidal activity (Jordan et al. 2012). Phenethyl alcohol, benzyl isothiocyanate, *β*-citronellol, geraniol, and nootkatone, (Dietrich et al. 2006, Tunon et al. 2006, Faraone, et al. 2019, Kim et al. 2021) are just some examples of compounds isolated from EO’s with repellent activity against ticks that provide opportunities for refinement and development of new marketable active ingredients.

Here, we describe the results of a semi-field trial using three modified monoterpenoid-derived compounds, ethyl perillyl carbonate (EPC), geranyl isovalerate (GI), and citronellyl cyclobutane carboxylate (CCC) where modifications were designed to reduce volatility. *In vitro* assays using treated filtered paper showed that two of the compounds (CCC and EPC), ester derivatives of compounds commonly found in plant oils, were highly repellent to *D. variabilis* (Wong et al. 2021). These monoterpene-based repellent molecules showed promise in the *in vitro* assays both in the short term and after five hours of aging (Wong et al. 2021). We tested the compounds for repellent activity against *I. scapularis* using treated sailcloth drags at an infested natural area in Wisconsin and compare the results with DEET and negative controls.

## Materials and Methods

### Repellents

The three compounds were synthesized in the laboratory but their simple ester chemistry is similar to naturally occurring compounds. One of the three, GI, is found in trace amounts in plants (Otomo et al. 1998). The compounds were obtained through Steglich esterification of monoterpenoid alcohols found in plant essential oils and one of a variety of small naturally occurring acids. The process is detailed in Wong (2021). Ethyl perillyl carbonate was derived from perillyl alcohol and ethyl chloroformate. Perillyl alcohol can be found in plants such as lavender and sage. Geranyl isovalerate was derived from geraniol and isovaleric acid. Geraniol is found in geraniums and roses and is a minor component of citronella oil. Citronellyl cyclobutane carboxylate was derived from citronellol, a monoterpenoid, and cyclobutane carboxylic acid. Citronellol is a minor component in citronella candles and found in plants such as lemongrass (*Cymbopogon citratus*) and lippia (*Lippia alba*). Compounds were used at the lowest concentration (0.16mg/cm^2^) where *in vitro* assays showed repellent activity against ticks that was similar to DEET.

### Semi-field Trial

#### Site Description

This study was conducted at the Big Eau Pleine County Park (BEP) in Marathon County (WGS84: 44.73912, -89.84111) where tick densities have historically been high during peak nymphal questing in the month of June and July. BEP is a mixed hardwood forest dominated by sugar maple (*Acer saccharum*) with an understory composed mainly of sedges, ferns and sugar maple saplings. At BEP, four plots were established based on similar topography and vegetation. Within each of the four plots, six subplots (40 × 10m; one subplot for each of four treatments and two negative controls) were established with 10 meter spacing between each subplot. Sampling occurred in June for the low concentration (LoC) trial and in July for the high concentration trial (HiC).

#### Repellent Field Assay

We used 24 dedicated tick drags (2840T: Bioquip Products, Rancho Dominguez, CA) for each of the 24 subplots and laundered drags after the LoC trial so that the same drags could be used for the HiC trial. DEET served as a positive control for this study. Negative controls included untreated (NEG) tick drags and acetone treatment only (ACE), as acetone was the carrier for all compounds. Repellents or acetone treatments were applied 15-20 minutes prior to the start of tick dragging and were applied to only one side of each tick drag using a 12oz spray bottle (Up&Up, Target, Minneapolis, MN) at a concentration of 0.16mg/cm^2^ (LoC) or 0.25mg/cm^2^ (HiC). At each plot, subplots and field technicians were randomly assigned to one of the six treatment groups using Microsoft Excel. Each subplot was dragged by a trained field technician and all subplots within a plot were dragged simultaneously. To assess the effectiveness of each compound against ticks, drags were placed on vegetation and technicians pulled them for 10 meters, repeating this 40 times for each 40 × 10m subplot. After each 10 meter sample the drag was hung on a shepherd’s hook (SKU:2779942, Menards, Eau Claire, WI) and checked for ticks. Any ticks initially found on the drag was identified to species and recorded. To further assess the repellent activity of the compounds against ticks, we observed all ticks (larvae, nymphs and adults) found on drags for up to 3 min. The time to detachment was recorded for each tick and all ticks that detached within 3 min were considered repelled. After the 3 min observation period, field technicians removed any remaining ticks from the drag and immediately returned them to the vegetation directly behind the collector.

#### Laundering of Drags

Once field technicians completed dragging of a subplot, the drag was placed in a black plastic garbage bag and sealed until they could be laundered. Drags were laundered between the LoC and HiC trials by soaking them in deionized water for 24-36 hours before being wrung out by hand and allowed to air dry for 3-5 days. All laundered drags were then tested for any unwanted residual repellency against ticks before being deployed again for the HiC trial. To evaluate residual repellent activity on drags, we randomly placed five field-collected nymphal *I. scapularis* ticks on each drag, held the drags vertically and counted the number of ticks that detached from each drag over a period of 5 min. Results showed that only one tick detached from a drag that was untreated in the previous LoC trial (data not shown). These results suggested that the process was sufficient to allow reuse of the drags for the HiC trial.

### Statistical Analysis

Evaluation of the compounds was done using R v4.0.3 (R Core Team 2021). Comparison of tick encounters (i.e. the proportion of drags positive for a nymphal tick) between repellent (DEET, CCC, GI, EPC) and control groups (NEG, ACE) was done using a Fisher’s Exact Test. A drag sample was considered positive if at least one nymphal tick was found after a10 meter drag. Comparison of the percentage of ticks that were repelled (detached before 3 min) was done using a Fisher’s Exact Test and the amount of time before detachment per tick was compared between repellent and control groups using a Kruskal-Wallis test with Bonferroni adjustment for multiple comparisons. All statistical analyses were considered significant at the alpha level of 0.05.

## Results

### Nymphs on Drags

We encountered a total of 314 nymphal *I. scapularis* on drags over the duration of the study (Table 1). We also encountered 257 larval *I. scapularis* but omitted them from further analysis because their small size and abundance made them difficult to count and track accurately in the field by a single technician. We encountered one adult *I. scapularis* on an ACE drag during the LoC trial and a single adult male *D. variabilis* was observed on a NEG drag in the HiC trial. More ticks were encountered during the trial period in June (LoC) compared to July (HiC; Table 1) which aligns with the phenological activity of *I. scapularis* nymphs in the area and is not a result of the treatment concentration.

**Table 1.**
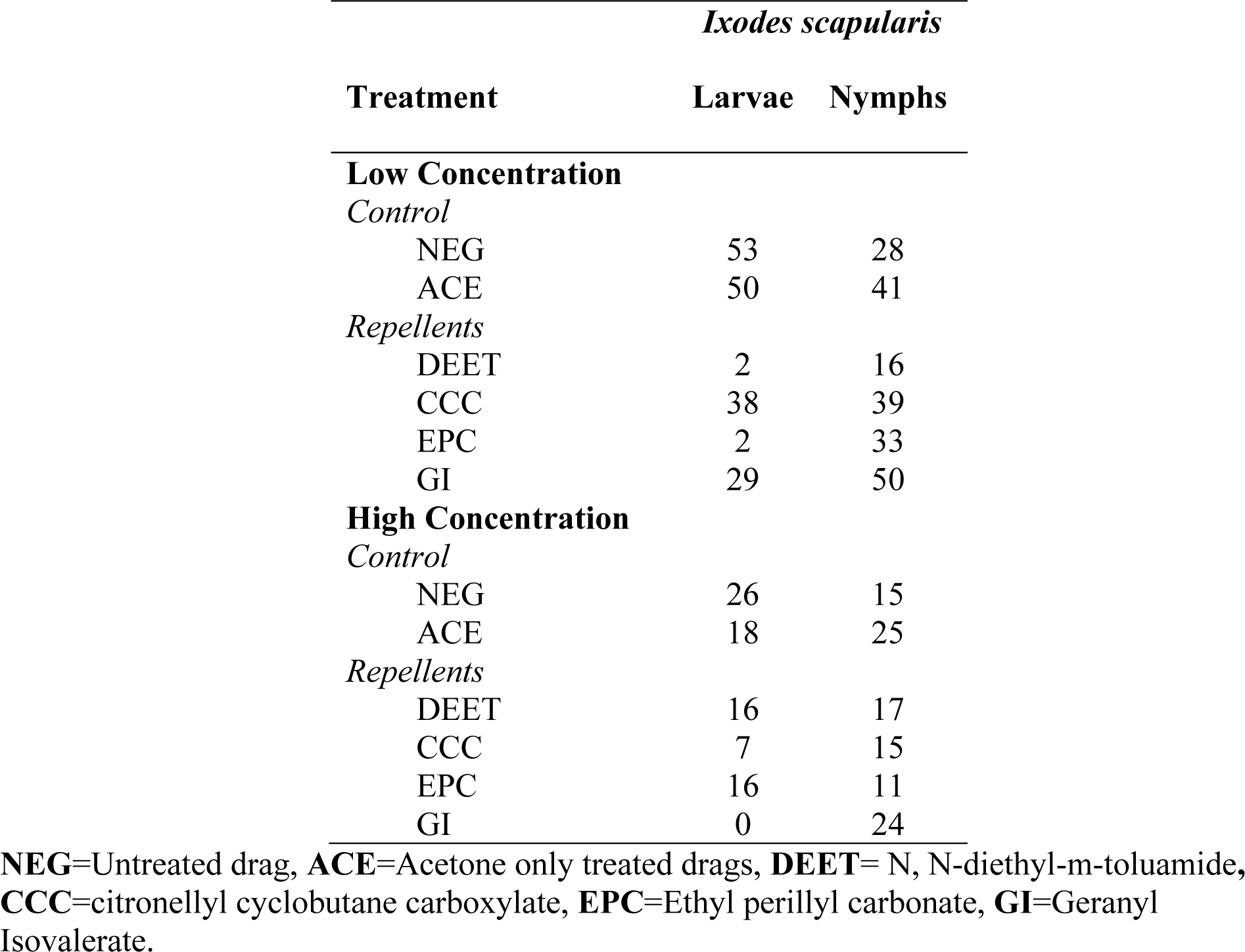
Summary of the total number of *I. scapularis* ticks encountered on drags during semi-field trials. Data shows the number of ticks collected from all subplots, a total of 160 drags (1,600 meters) for each treatment.

No treatment repelled all ticks and there was significant variation even between the two negative control treatments (Table 1, Figure 1). None of the repellent treatments resulted in significantly fewer tick encounters in comparisons with both of the negative controls. The number of 10 meter drag samples (N=160 for each treatment) positive for at least a single nymphal *I. scapularis* tick was significantly lower for DEET (15 drags) when compared to ACE (36 drags), CCC (30 drags) and GI (45 drags) in the LoC trial (p<0.02; Figure 1) but was not different from NEG. There were no significant differences (p>0.1) in the number of drags positive for nymphs between NEG, ACE, CCC and EPC. The number of positive drags was significantly higher (p<0.01) for the GI treatment compared to NEG (25 drags) and EPC (25 drags).

**Figure 1.**
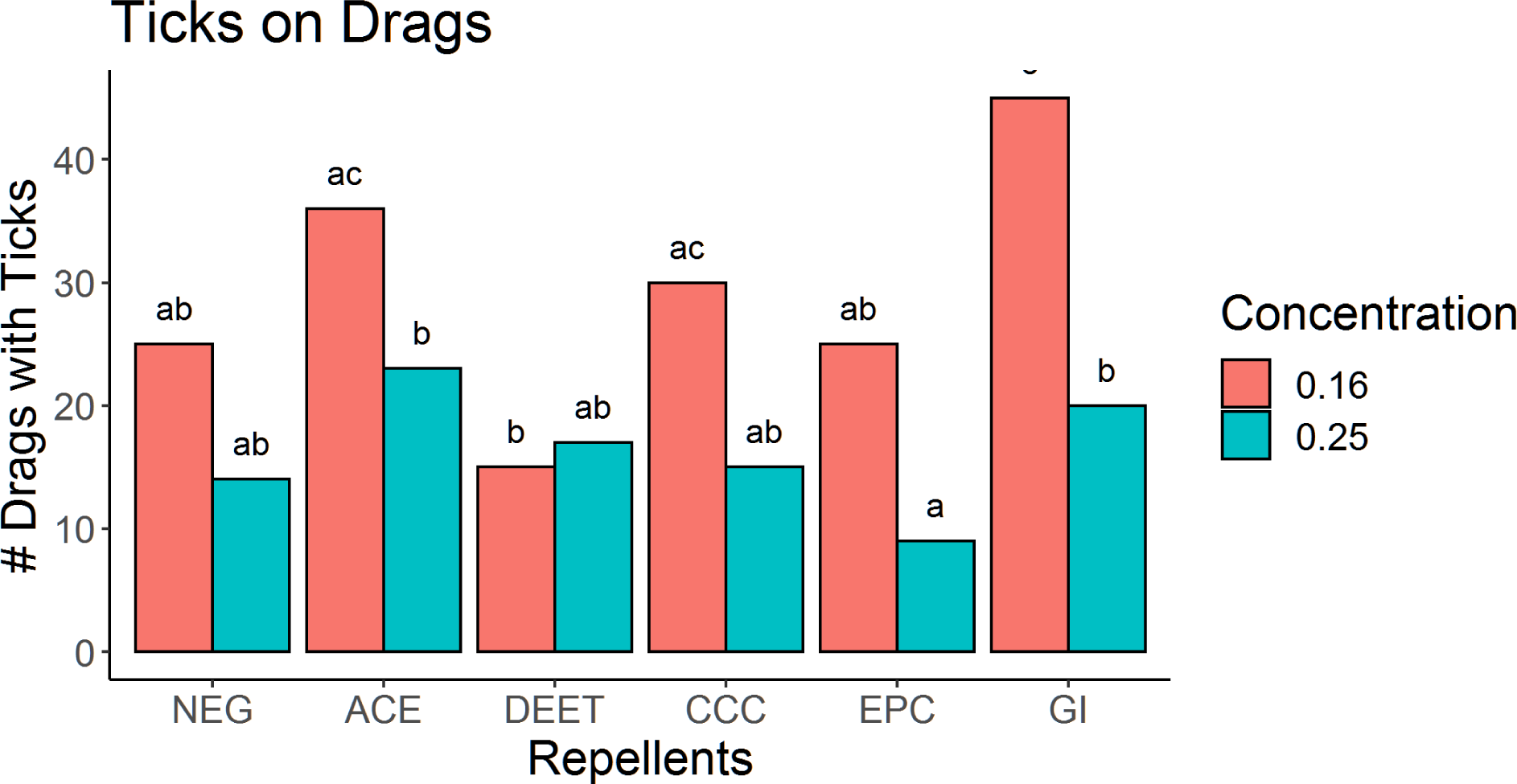
Tick encountered on drags defined as the number of drags with at least one nymphal *I. scapularis* after 10 meters of dragging. Letters denote statistical significance (Fisher’s Exact Test) within the same concentration trial. **NEG**=Untreated drag, **ACE**=Acetone only treated drags, **DEET**= N, N-diethyl-m-toluamide, **CCC**=citronellyl cyclobutane carboxylate, **EPC**=Ethyl perillyl carbonate, **GI**=Geranyl Isovalerate

In the HiC trial, we saw no significant differences (p>0.1) in the number of drags with at least a single nymphal *I. scapularis* between NEG (14 drags), DEET (17 drags), CCC (15 drags) and EPC (9 drags). The EPC treatment was associated with the lowest number of positive drags (9 drags) but this was significantly lower only when compared to ACE (23 drags; p=0.014) or GI (20 drags; p=0.049).

### Ticks Repelled During Observation Period

Very few nymphal ticks detached from the negative control drags during the 3 min observation period. Detachment from NEG and ACE drags was significantly lower (p<0.001) compared to all other repellents in both the LoC and HiC trials (Figure 2). By contrast, the percentage of nymphal *I. scapularis* ticks that detached within the observation period was high on drags treated with repellents. For example, we observed detachment of 87.5% and 69.7% of nymphs on DEET and EPC treated drags, respectively during the LoC trial (Figure 2). The percentage of ticks repelled by DEET was significantly higher than for CCC or GI but not EPC. However, detachment from EPC-treated drags was not significantly different from drags treated with CCC (56.4%) or GI (48%; p>0.07) or from DEET. The patterns were similarly distinct between negative controls and repellents during the HiC trial and there were no significant differences (p>0.1) in the percentage of detached ticks between the four repellents (Figure 2).

**Figure 2.**
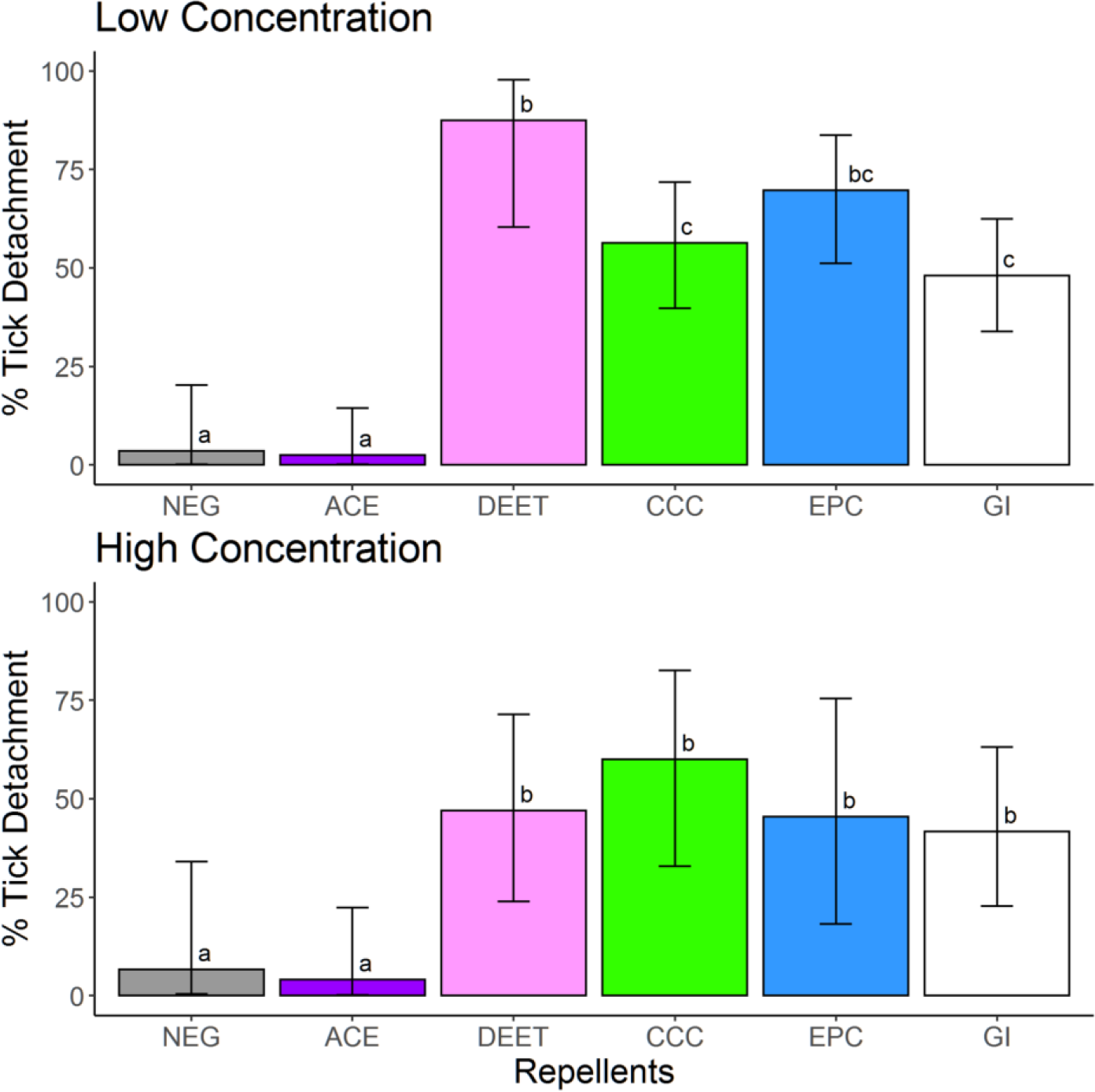
Percentage of nymphal *I. scapularis* ticks that detached from drags within a 3 minute observation period. Low concentration is 0.16mg/cm^2^ and high concentration is 0.25mg/cm^2^ of repellent or acetone only applied to each drag. Letters denote statistical significance (Fisher’s Exact Test) and bars represent the 95% confidence intervals for each proportion. **NEG**=Untreated drag, **ACE**=Acetone only treated drags, **DEET**= N, N-diethyl-m-toluamide, **CCC**=citronellyl cyclobutane carboxylate, **EPC**=Ethyl perillyl carbonate, **GI**=Geranyl Isovalerate.

### Total Attachment Time

In general, the median nymphal detachment time was lowest on drags treated with DEET and EPC during both LoC and HiC trials but this was not significantly different when compared to the other repellents (p>0.1; Figure 3). NEG and ACE were excluded from time analysis because only one nymphal *I. scapularis* tick detached from the negative control drags during the LoC and HiC trials.

**Figure 3.**
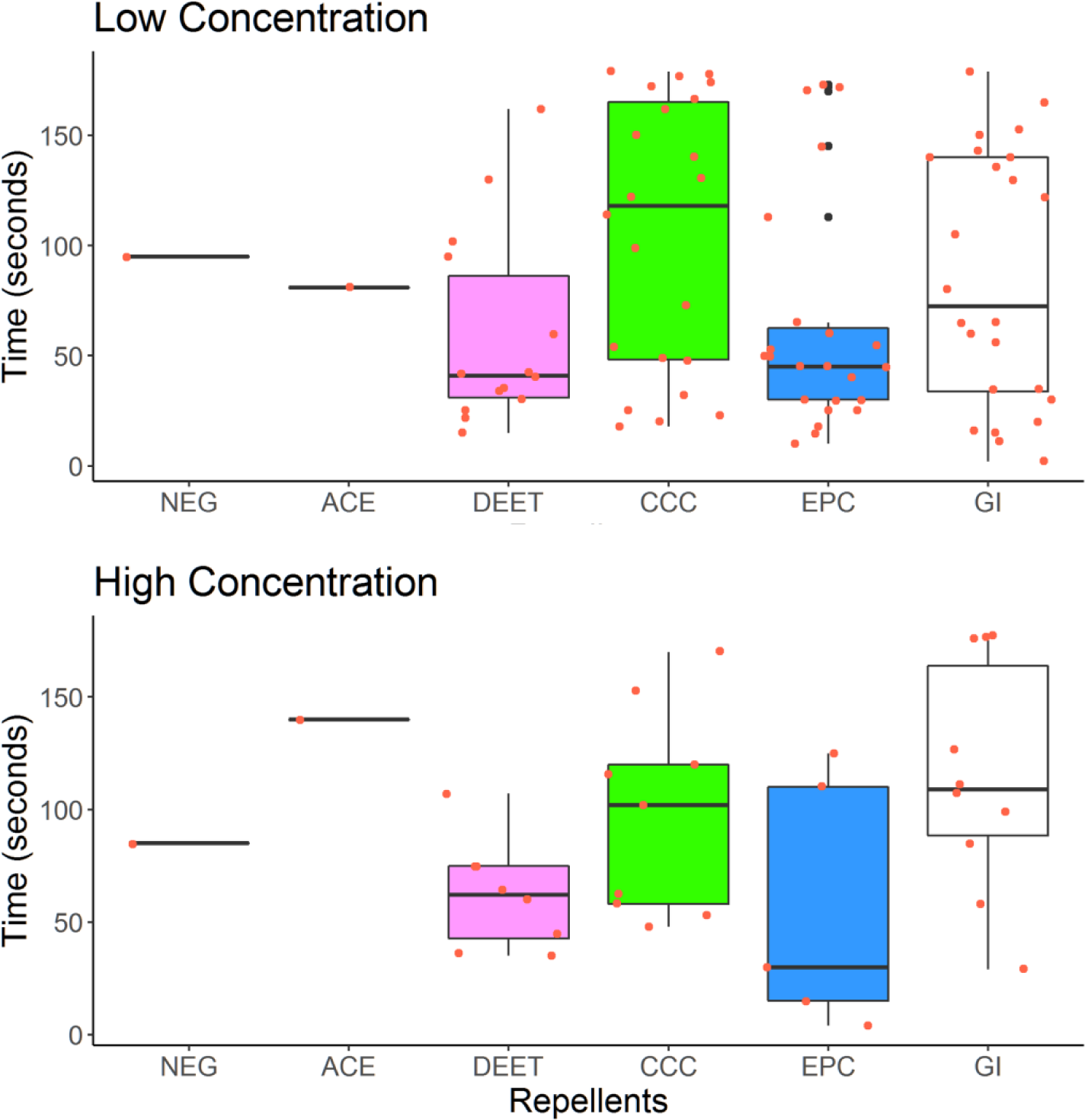
Median detachment time for nymphal *I. scapularis* during a 3 minute observation period. Low concentration is 0.16mg/cm^2^ and high concentration is 0.25mg/cm^2^ of repellent or acetone only applied to each drag. **NEG**=Untreated drag, **ACE**=Acetone only treated drags, **DEET**= N, N-diethyl-m-toluamide, **CCC**=citronellyl cyclobutane carboxylate, **EPC**=Ethyl perillyl carbonate, **GI**=Geranyl Isovalerate.

## Discussion

Many botanically derived repellents have been described based on laboratory trials but few of these have been validated using real-world conditions. Ticks used in the laboratory studies are not host-seeking and behave differently from ticks that are actively “questing”. We sought to build on promising lab results by testing the compounds against active, host-seeking nymphs in a natural setting and were able to confirm strong repellency against nymphal *I. scapularis*. In addition, treating drag fabric proved to be a useful method that enabled rapid evaluations without costly and time-consuming procedures involved in repellent research that requires human subjects.

We report that three recently derived compounds (CCC, EPC and GI) had repellent activity against *I. scapularis* ticks similar to the current gold standard repellent for all comparisons, DEET. For all four repellent treatments, nymphs initially attached to treated drags but were quickly repelled and detached while nearly all nymphs on untreated or acetone-treated drags remained attached. The percentage of nymphs that detached for the three test compounds ranged from 48% to 87% in LoC and from 42% to 60% in HiC (Figure 2). The higher concentration of compounds in HiC did not result in higher detachment. It is likely that the maximum effective concentration-repellency of DEET and the three compounds may have been reached at a concentration of 0.16mg/cm^2^ and higher concentrations would not yield additional repellency benefits (Pages et al. 2014, Diaz et al. 2016, Meng et al. 2016). Regardless, our study shows that the three compounds are as effective as DEET in preventing tick encounters and detachment.

A limitation in our study is that we could not assure that drags were uniformly covered with the test compounds. Drags were sprayed with 145mL of each compound in acetone until the drag was visibly wet. Unfortunately, there was no way to ensure complete coverage of drags in the field and incomplete coverage could certainly have impacted median detachment times. Median detachment time for ticks was not significantly different between DEET and the three compounds, but a range of detachment times were observed (Figure 3). Despite the variation in detachment times within the three minute observation period, all three compounds were successful at causing ticks to detach from drags once attached. Future studies should include more uniform treatment of fabrics as well as longer observation periods to fully assess the effects.

As a repellent, long lasting protection is a vital component for success. Short-lasting, volatile compounds have limited impacts and the need for frequent reapplication can reduce protection against tick bites (Soutar et al. 2019). Due to time and personnel constraints, we could not test the effective duration of the three compounds. However, we surmise that, at a minimum, the effective duration would be equivalent to the time needed to complete the dragging of each subplot. Field technicians spent on average, 1-2 hours dragging each treatment subplot and we did not detect any notable difference in the frequency of tick encounters or detachment times for drags completed during the last quarter section of each subplot (data not shown). These observations support an effective duration of at least 1-2 hours. Laboratory assays conducted by Wong et al. (2021) showed that EPC treated filtered paper that was aged for 5 hours repelled greater than 40% of adult *I. scapularis*. Unmodified monoterpenoid alcohols can be repellent, but often lack longevity due to high volatility. The semi-synthetic esters used in this study were developed to enhance the effective duration of repellent activity (Norris and Coats 2017, Wong et al. 2021) but further studies will need to be conducted to fully examine the longevity of the compounds under field conditions.

In summary, our study showed that three recently derived compounds were effective against *I. scapularis* nymphs in a semi-field study. Additional research on the effective duration and *in vivo* assays using humans will be necessary to assess the full potential and protection that these compounds may provide. Our hope is that these new compounds may provide the public with a “green” repellent option in various formulations and concentrations.

## Acknowledgements

We thank the fellows from the Midwest Center of Excellence for Vector-borne Disease. We thank Jamie Polley and members of the Wausau and Marathon county parks, recreation and forestry department for permission to conduct this study at the Big Eau Pleine county park. This work was supported by Cooperative Agreement Number U01CK000505, funded by the Centers for Disease Control and Prevention. Its contents are solely the responsibility of the authors and do not necessarily represent the official views of the Centers for Disease Control and Prevention.

